# An algorithm-centric Monte Carlo method to empirically quantify motion type estimation uncertainty in single-particle tracking

**DOI:** 10.1101/379255

**Authors:** Alessandro Rigano, Vanni Galli, Krzysztof Gonciarz, Ivo F. Sbalzarini, Strambio-De-Castillia Caterina

## Abstract

Quantitative analysis of microscopy images is ideally suited for understanding the functional biological correlates of individual molecular species identified by one of the several available “omics” techniques. Due to advances in fluorescent labeling, microscopy engineering and image processing, it is now possible to routinely observe and quantitatively analyze at high temporal and spatial resolution the real-time behavior of thousands of individual cellular structures as they perform their functional task inside living systems. Despite the central role of microscopic imaging in modern biology, unbiased inference, valid interpretation, scientific reproducibility and results dissemination are hampered by the still prevalent need for subjective interpretation of image data and by the limited attention given to the quantitative assessment and reporting of the error associated with each measurement or calculation, and on its effect on downstream analysis steps (i.e., error propagation). One of the mainstays of bioimage analysis is represented by single-particle tracking (SPT)^1–5^, which coupled with the mathematical analysis of trajectories and with the interpretative modelling of motion modalities, is of key importance for the quantitative understanding of the heterogeneous intracellular dynamic behavior of fluorescently-labeled individual cellular structures, vesicles, virions and single-molecules. Despite substantial advances, the evaluation of analytical error propagation through SPT and motion analysis pipelines is absent from most available tools ^6^. This severely hinders the critical evaluation, comparison, reproducibility and integration of results emerging from different laboratories, at different times, under different experimental conditions and using different model systems. Here we describe a novel, algorithmic-centric, Monte Carlo method to assess the effect of experimental parameters such as signal to noise ratio (SNR), particle detection error, trajectory length, and the diffusivity characteristics of the moving particle on the uncertainty associated with motion type classification The method is easily extensible to a wide variety of SPT algorithms, is made widely available via its implementation in our Open Microscopy Environment inteGrated Analysis (OMEGA) software tool for the management and analysis of tracking data ^7^, and forms an integral part of our Minimum Information About Particle Tracking Experiments (MIAPTE) data model ^8^.

## Background

Time series of optical images acquired from living cells are the starting point for the analysis of the intracellular dynamics and interactions of myriads of heterogeneous cellular features ^9–26^. Single-particle Tracking (SPT) experiments entail three distinct steps (Figure 1): particle detection and tracking, trajectory analysis ^5,11,27–38^, and physical interpretation through modeling ^17,39–41^. The first step consists of detecting diffraction-limited objects (i.e., fluorescently labeled vesicles, organelles, single molecules or viral particles) in a digital image time series, and of linking these detections (i.e., feature points) over time with the ultimate purpose of reconstructing their trajectories as they move within living organisms. The end result of this first step is therefore a sequence of spatial coordinates indicating the position of each target particle at each time point. After localization and linking, both local and global descriptive metrics (e.g., direction, displacement, velocity, isotropy and diffusion constants) have to be computed from individual trajectories in order to provide a biological interpretation to the movement behavior and molecular interactions of the biological structure under study (reviewed in ^11,42^). Last but not least, the “interpretation problem” ^6^ consists in the inference of motion models (i.e., confinement, normal diffusion, drift, and directed motion) and their parameters from such descriptive trajectory metrics with the ultimate goal of understanding the underlying biological significance of changes in direction, velocity, and freedom of movement (i.e., interactions with an heterogeneous environment might lead to frequent changes in diffusivity and even result in temporary stoppages).

**Figure 1:**
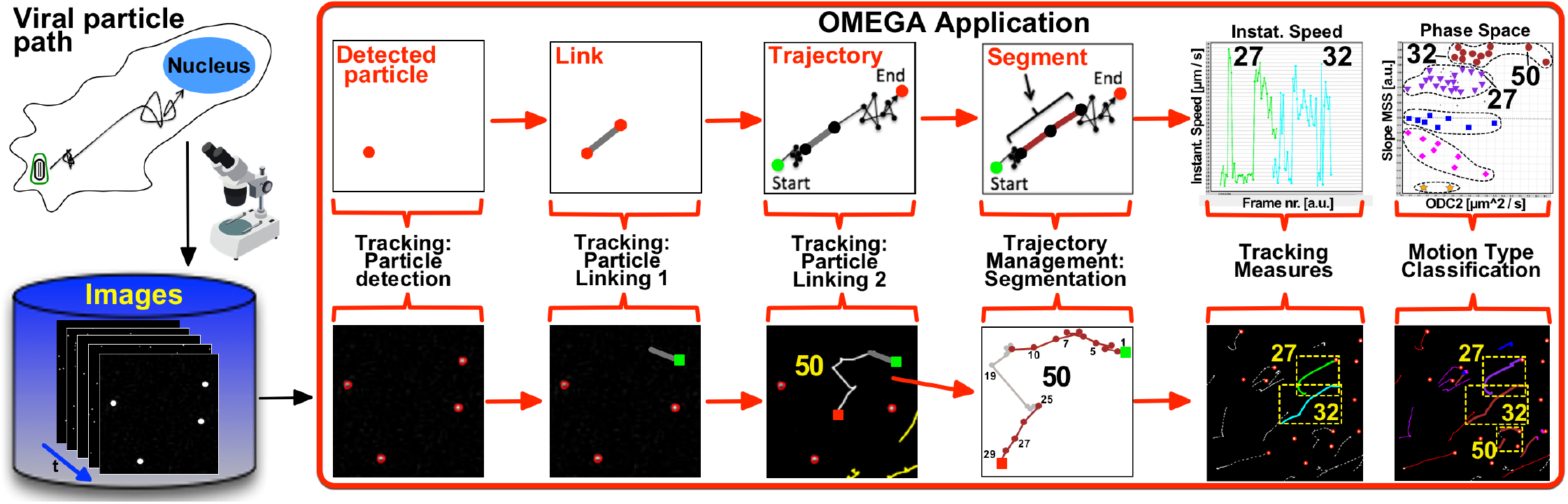
Typical particle tracking and motion analysis workflow in OMEGA. Schematic diagram depicting the system context in which OMEGA operates and the workflow required for the estimation of the sub-cellular trajectories followed by diffraction-limited intra-cellular viral particles and the computation of biologically meaningful measures from particles coordinates. After acquisition using available microscopes, images are loaded onto the OMERO image and metadata repository (*blue*), and subsequently subjected to single particle tracking (SPT) using the two modular Particle Detection and the Particle Linking plugins in OMEGA. All trajectories which appeared uniform were assigned the color corresponding to the predicted motion type depending on the observed slope of the MSS curve (*grey*, unassigned; *yellow*, confined; fuchsia, sub-diffusive; *blue*, diffusive; *purple*, super-diffusive; *maroon*, directed). As needed individual trajectories can be subdivided in uniform segments using the interactive OMEGA Trajectory Segmentation plugin. In the example, trajectory nr. 50 was subdivided in three segments two of which were assigned the Directed motion type (*maroon*) and the third one was left un-assigned (*grey*). Resulting trajectories an segments were then subjected to motion analysis using the Velocity Tracking Measures and the Diffusivity Tracking Measures plugins. Instantaneous Speed results for trajectory nr. **27** and **32** and ODC vs. SMSS Phase space results for all trajectories are displayed. The position of spots representing trajectory nr. **27, 32** and **50** are indicated.

Because of their size, intracellular vesicles, single molecule, virions and other diffraction-limited objects behave like Brownian particles, which when unperturbed are expected to freely diffuse ^43^. Under these general conditions, deviations from normal diffusion can result from interactions that alter the rate or direction of motion. In the case of viral particles for example ^2,3,11,21,44–46^, interaction with motor proteins might result in directed motion along microtubules. Alternatively, interaction with a relatively immobile cellular structures such as membrane rafts at the cellular periphery or nuclear pore complexes at the nuclear envelope, might result in the transient confinement of viral particles to restricted zones. As a corollary, free diffusion is one of the most common models employed to interpret experimental particle tracking data. It is therefore of fundamental importance to understand how accurately free diffusion or other motion types can be estimated from finite length trajectories with determinate localization accuracy and precision, especially in the frequent cases where it is necessary to assess the statistical significance of apparent differences between distinct segments of the same trajectories, different trajectories within the same sample or between results of experiments performed under varying conditions.

Diffusion is a random process and its study is based on the statistical analysis of subsequent recordings of the object’s position as it moves in space across time. Assuming that all motion processes, irrespective or their diffusive characteristics can be described in terms of *probability*, it is therefore possible to classify the dynamic behavior of individual particles using the same method irrespective of their motion properties. A basic metric describing microscopic dynamics is the observed diffusion constant (ODC), which provides meaningful information regarding the quantity of displacement, and which might remain constant in an isotropic medium or vary in space and time in complex environments. Specifically, different motion regimens can be classified by computing ODC and by combining it with a measure of freedom of motion, such as the variation of Mean Square Displacement (MSD) over time ^5^ or the Hurst exponent (henceforth referred to as the Slope of the Moment Scaling Spectrum or SMSS) ^47,48^, thereby representing individual trajectories as points in phase space ^40,49^. In addition to representing a massive data reduction, this approach facilitates the classification of the mobility characteristics of multiple particles all at once without arbitrary selection.

While global metrics such as ODC and SMSS offer powerful analytic tools, it is important to emphasize SPT can be misleading if used incorrectly. Despite considerable advances in trajectory analysis, uncertainty estimation and reporting has so far received too little attention. In particular, the consequence of finite localization accuracy, precision and observation times (i.e., trajectory length) on the estimation of metrics such as ODC, MSD and MSS has been scarcely addressed ^6^. Error associated with each step of the particle tracking workflow is not routinely estimated and its effect on downstream analysis steps are often not taken into account. This can lead to significant problems with interpretation, reproducibility and re-use of analysis results. Whatmore, because the clear definition and interpretation of uncertainty metrics is not trivial, even when methods to estimate particle tracking error are developed they are typically not universally adopted or easy to use, compounding the problem of results dissemination.

In general, trajectory motion analysis error might derive from one of three possible sources: 1) the presence of raw image noise is typically propagated in the analysis pipelines and leads to often amplified uncertainty in the results; 2) algorithms might produce inherently biased or suboptimal results; and 3) inadequacies in the *ex ante* interpretative models of the system under study may result in uncertain results (a.k.a. model misspecification error). The problem is exacerbated by the absence of ground truth, the inadequacy of synthetic benchmarks and lacking theoretical frameworks to guarantee algorithm performance ^49^. Promising approaches have emerged especially for what concerns the generation of highly curated collections of error-free, real-life ^50^ or synthetic ^32,51^ benchmarks on which to compare the performance of algorithms and provide error assessment. On the other end the empirical analysis of error propagation for specific algorithms ^52–54^, has been so far quite rare in image analysis in general and particle tracking in particular.

To obviate this difficulty, the algorithm-centric empirical method described here utilizes Monte Carlo simulation to generate sets of ideal trajectories, utilizes artificial images of point emitters in the presence of known amount of noise to empirically derive distributions of observed localization errors for each algorithm under study, simulates the effect of localization noise on ideal trajectories by offsetting the position of each detection with randomly sampled error values from such pre-computed distributions, and estimates the probability density of observed results to predict the error associated with potential results. As a proof of concept the approach concentrates on the estimation of error associated with the MOSAICsuite Feature Point Tracking algorithm ^49,55,56^ and on the computation of ODC and SMSS as implemented in the recently released Open Microscopy Environment inteGrated Analysis (OMEGA) software platform (Figure 1) ^7^. However the method is designed to be easily extendable to other particle detectors and to be used for other global trajectory measures.

## Outline

The numerical procedure described here empirically assesses the estimation accuracy for ODC and SMSS obtainable with a specific particle detection algorithm, given the observed localization error of each detected point feature in a given signal to noise ratio (SNR) condition, the trajectory length and the motion type characteristics (i.e., ODC and SMSS) of the moving particle. While the procedure can be applied to any particle detection/localization algorithm, results presented here refer to the MOSAICsuite Feature Point Detection (FPD) algorithm ^55,56^ as implemented in the Open Microscopy Environment inteGrated Analysis (OMEGA) particle tracking software ^7^ (Figure 1). In general, the procedure can be subdivided in three steps:

1. Estimation of the particle detection/localization error associated with a given algorithm and under a set of representative SNR conditions. This procedure is conducted for each algorithm that one wishes to utilize and the resulting inaccuracy statistics are then stored either locally or in shared databases for future use by the community (Figure 2).
2. Evaluation of the effect of the inaccuracy distribution associated with each particle detection algorithm and SNR value on the estimation of ODC and SMSS given test datasets composed of artificial trajectories of known trajectory length, SMSS and ODC (Figure 3).
3. Implementation of the method used to estimate the motion type classification error associated with each ODC and SMSS calculation on the basis of previously stored empirical uncertainties distributions and of the desired confidence interval (Figure 5).

**Figure 2:**
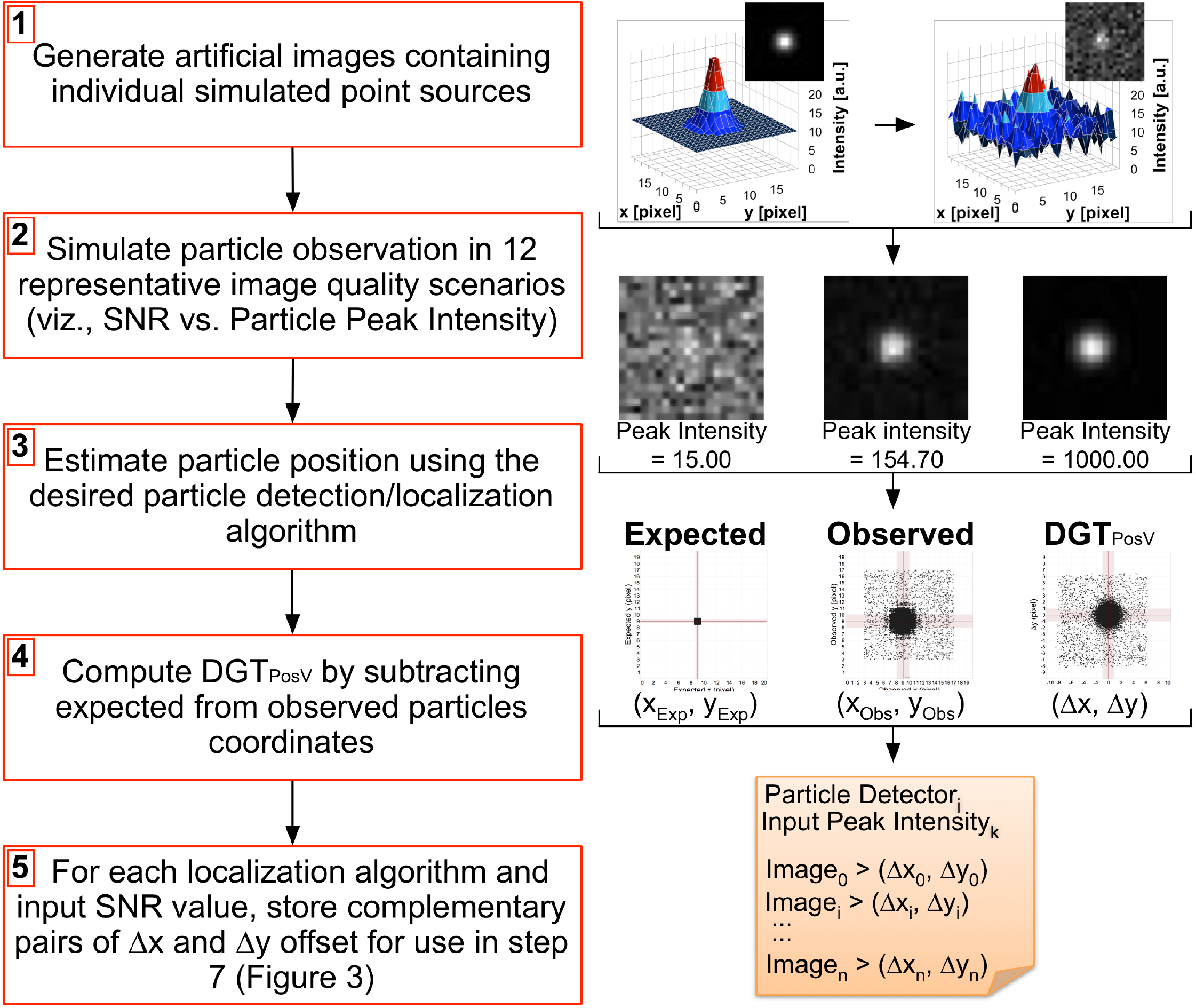
Determination of the localization accuracy and precision associated with examined particle detection/localization algorithms. The localization accuracy (i.e., bias = arithmetic mean error) and precision (i.e., sigma = standard deviation of the error) of each examined particle detection/localization algorithm is empirically estimated using a set of artificial images (see Supplemental Material) using the indicated steps: **(1)** Sets of 20 x 20 pixel images are generated each containing one single simulated point emitter placed at a known random position (i.e., ground truth) within the central pixel. Microscopic observation is simulated by sampling a Gaussian of standard deviation σ = 1 pixel and mean μ = particle peak intensity (P_PI_), cantered at each ground truth point position. **(2)** In order to simulate different image qualities (viz., signal to noise ratio = SNR), point sources are assigned twelve P_PI_ values (Supplemental Table I), and Poisson-distributed Shot noise associated with CCD camera image acquisition, is modelled by replacing the peak intensity I of each pixel with a number sampled at random from a Poisson distribution of mean value μ = λ = I. In all cases background value is kept constant at b = 10. **(3)** The Cartesian Positional Vector (PosV) of each simulated point source present in the benchmarking test sets is determined for each examined P_PI_ value and particle detection/localization algorithm. **(4)** Distributions of observed Distances from Ground Truth (DGT) of PosV (*DGT_PosV_*) are computed by subtracting *Expected* (i.e., ground truth; input) from *Observed* (i.e., output) PosVs, plotted on scatter plots alongside *Expected* (i.e., ground truth) and *Observed* point position coordinates (Supplemental Figure 2), and **(5)** stored for later use (Figure 3, Step 7).

**Figure 3:**
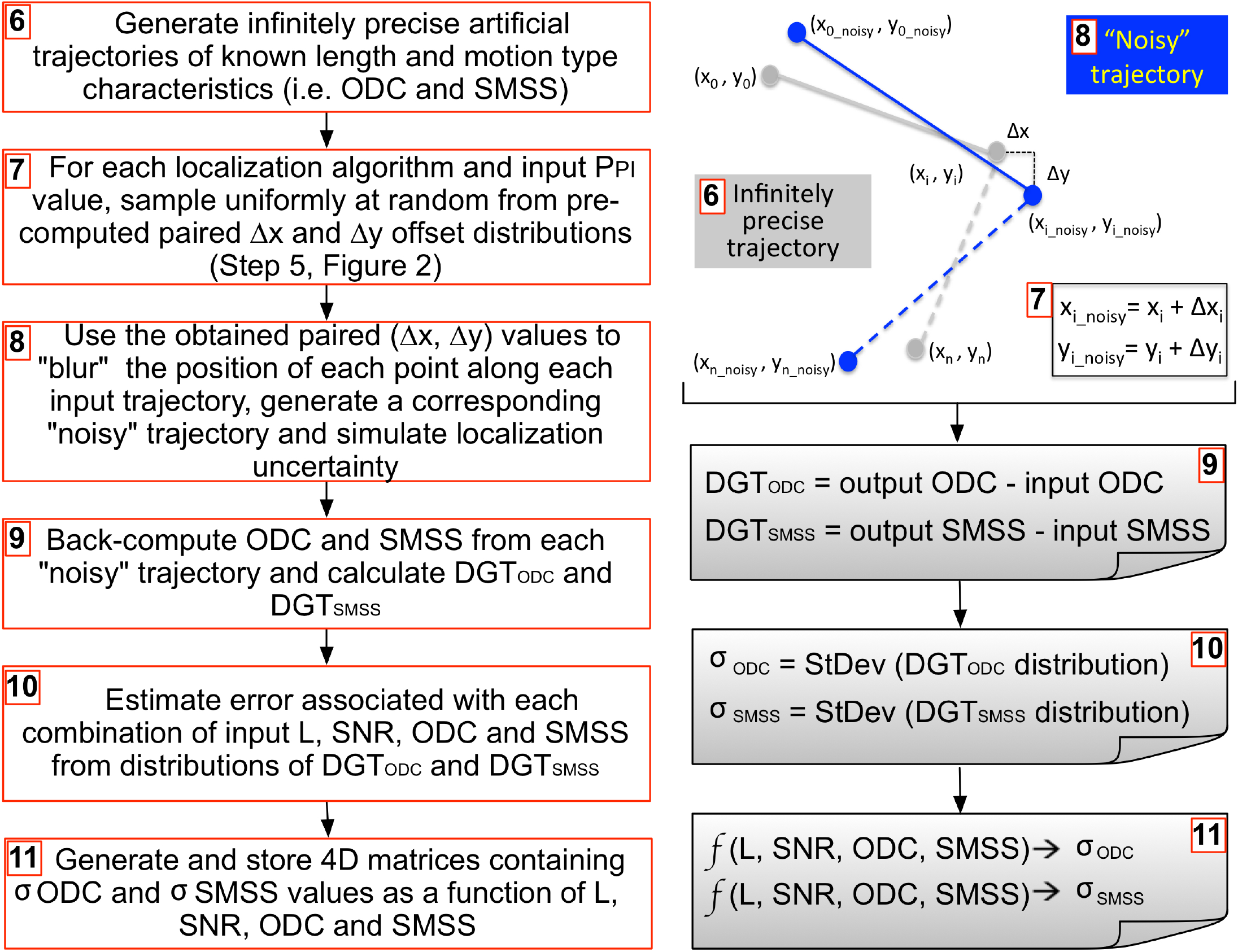
Empirical estimation of the uncertainty associated with key diffusivity measures based on observed particle localization uncertainty. Observed diffusion constant (ODC) and the slope of the moment scaling spectrum (SMSS; Ferrari *et al.*, 2001), are used in OMEGA to determine the diffusive characteristics of diffraction-limited particles of interest. Steps utilized to empirically asses the uncertainty associated with ODC and SMSS estimation are as follows: **(6)** Infinitely precise trajectories of known input length, ODC and SMSS are artificially generated using our custom-made *artificialTrajectories2* Matlab algorithm (Helmuth *et al*, 2007). **(7)** For each examined particle detection/localization algorithm and each modeled particle peak intensity (PPI) value (Supplemental Table I), a pre-computed data set of paired Δx and Δy offset distributions (Figure 2, Step 5), is sampled uniformly at random. **(8)** The obtained paired set of Δx and Δy offsets are added to the coordinates of all points along each trajectory to obtain a set of corresponding “noisy” trajectories. **(9)** Output ODC and SMSS values are then back-computed from each “noisy” trajectory and corresponding Distance from Ground Truth (DGT) distributions are obtained for each measure by subtracting expected (*input*) from observed (*output*) values. **(10)** For each measure, uncertainties associated with each combination of input parameters (i.e., L, SNR, ODC, SMSS) are estimated as the standard deviation of the corresponding DGT distribution. **(11)** Uncertainties are stored for later use in four-dimensional (4D) matrices with L, SNR, ODC, SMSS indexes (Figure 5, Step 6).

## Results

### Estimation of the error associated with a given particle detection/localization algorithm

Because the trueness of a given spot detector algorithm cannot be estimated unless the correct position of each particle under observation is completely known, the performance of particle detection/localization algorithms is generally assessed using artificial images (Figure 2). Specifically, in order to empirically assess and record the localization accuracy and precision of the OMEGA MOSAICsuite FPD plugin ^7,49,55,56^, individual simulated single point fluorescent sources were placed at random known sub-pixel positions (i.e., ground truth) within the central pixel of 20 x 20 pixel synthetic planes as described (Materials and Methods; Supplemental Figure 1) ^7,49^. To simulate different image qualities, images were generated to reflect twelve different input peak intensity and corresponding SNR values (Supplemental Table I) and the stationary feature points present in each synthetic plane were then localized using the OMEGA plugin mentioned above (Supplemental Figure 2). Finally, distributions of localization errors associated with each detected particle were computed to evaluate tracking quality (Supplemental Figure 3). In generalized terms, for each examined particle detection/localization algorithm under study and input SNR value, expected Cartesian positional vector (PosV) are compared with observed PosV, and distributions of observed DGT of the PosV (DGT_PosV_) are computed and stored as follows:

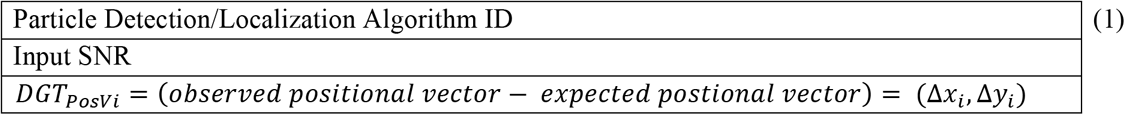

If needed these distributions can be used to compute localization bias and standard deviation (i.e., sigma which in turn are used as measures of accuracy and precision respectively. Thus:

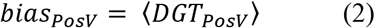

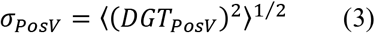

Where: 〈 〉 denote ensemble averages over independent trials; bias_PosV_ equals the average of all observed DGT_PosV_ values; and standard deviation. As shown in Supplemental Figure 3 and as expected, the values of both bias and sigma decreased with image quality and converged toward zero.

### Empirical evaluation of the uncertainty associated with the estimation of ODC and SMSS

The method presented here is based on the *in silico* Monte Carlo simulation of artificial trajectories, whose true position with respect to the imaging system, rate of displacement (i.e., ODC), freedom of movement (i.e., SMSS) and length (i.e., L) and are fully known (Figure 3). After modeling the effect of positional error on these “ground truth” trajectories under different particle peak intensity (P_PI_) and associated local SNR conditions, ODC and SMSS were back-computed from the resulting “noisy” trajectories and the comparison between input and output values was used to estimate the quantity of error associated with motion type estimation as a function of expected motion characteristics (i.e., ODC and SMSS), motion duration (i.e., L) and local SNR.

Specifically, in order to empirically assess ODC and SMSS estimation error, 1936 test sets (16 x L; 11 x ODC; 11 x SMSS; Supplemental Table I) each containing 1000 artificial two-dimensional trajectories were generated using our *artificialTrajectory2* MatLab function ^7,49^, and assuming the following physical dimensions: 1 pixel/μm and 1 frame/sec (see Materials and Methods). Such trajectories were then subjected to positional blurring as described in Material and Methods to model the effect of 12 representative input P_PI_, of associated local SNR values (Supplemental Table I) and of particle detection/localization as determined by using the MOSAICsuite FPD algorithm available as an OMEGA plugin ^7^. Thus, the OMEGA Diffusivity Tracking Measures (DTM) plugin ^7^ was used to back compute observed values of ODC and SMSS from the resulting 23232 (i.e., 1936 x 12 local SNR_input_ values) test sets of “noisy” trajectories ^7^, employing L/3, L/5 and L/10 calculation-window sizes ^7^. Finally, distributions of DGT for both ODC (i.e., DGT_ODC_) and for SMSS (i.e., DGT_SMSS_) were calculated for each combination of input parameter values as follows:

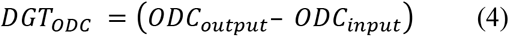

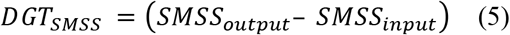

Standard deviations obtained from these distributions of DGT_ODC_ and DGT_SMSS_ were utilized as a measure of estimation uncertainty, saved for each particle detection/localization algorithm to be considered, and stored as ready-to-use four-dimensional uncertainty matrixes as follows (Figure 4):

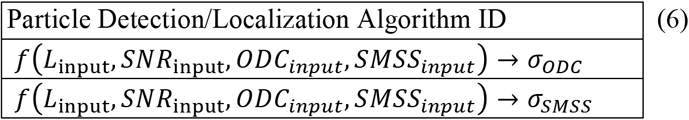

**Figure 4:**
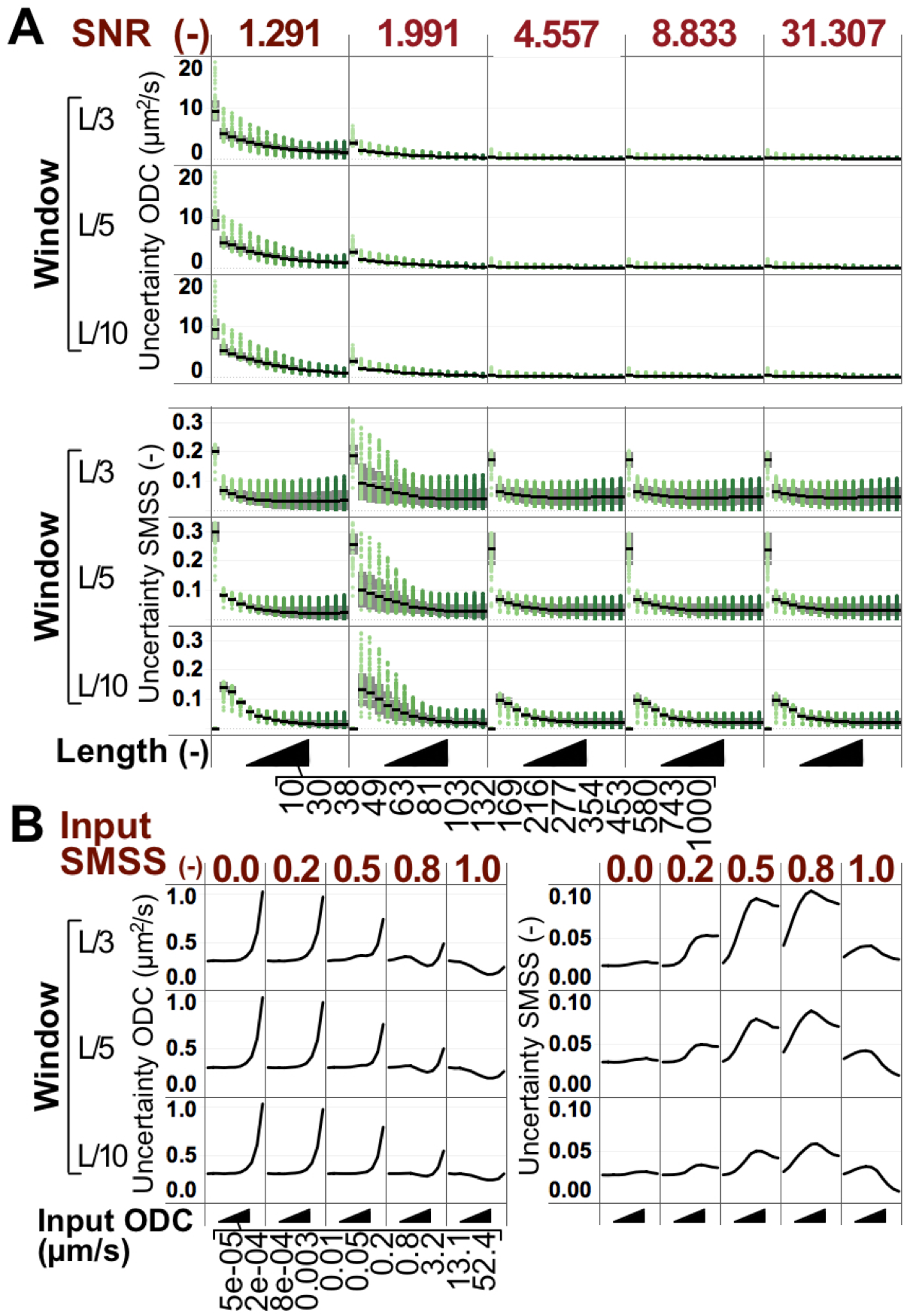
ODC and SMSS estimation uncertainty depend on SNR, trajectory length and on the diffusivity behavior of moving particles. In order to compute simulated ODC and SMSS estimation uncertainties, 1936 test sets representing 16x L, 11x ODC and 11x SMSS input values (Supplemental Table I) each containing 1000 artificial trajectories, were generated using *artificialTrjectories2* (Helmuth *et al.*, 2007), and subjected to position blurring on the basis of localization offsets observed under 12 representative SNR conditions and using the MOSAICsuite Feature Point Detection (FPD) algorithm. The OMEGA Diffusivity Tracking Measures (DTM) plugin was employed to back compute observed ODC and SMSS values from these “noisy” trajectories, using computation windows equal to L/3 and L/5 and L/10 as indicated on the left of each plot. Distributions of Distances from Ground Truth (DGT) were calculated for each combination of input parameter values, and standard deviations from these distributions were used as a measure of uncertainty for each measurement (see Figure 3). **(A)** ODC (Top) and SMSS (*Bottom*) estimation uncertainty values are presented here as scatter distributions of individual values and plotted as a function of input SNR (vertical columns as indicated by red labels) and L (x axis as indicated). As expected in both cases uncertainty diminishes with increasing SNR and trajectory length Black lines indicate average values and grey area indicated the +/-1 standard deviation interval around the mean. **(B)** Average values for both ODC (*Left*) and SMSS (*Right*) estimation uncertainty distributions computed as above are displayed in relation to input SMSS (vertical columns as indicated by red labels) and ODC (x axis as indicated) input values. Estimation of ODC was observed to be more difficult for extreme ODC input values and easier for higher input SMSS values. Estimation of SMSS was observed to be easier at the extremes of both input SMSS and ODC. ODC values were calculated in μm^2/s assuming 1 pixel/μm and 1 frame/s.

Where L_input_, SNR_input_, ODC_input_, and SMSS_input_ are the values of trajectory length, SNR, ODC and SMSS that were used as input during trajectory generation. As expected, when ODC estimation uncertainty values (see Eq. 4, 5 and 6) were plotted as a function of trajectory length and local SNR, they were observed to diminish with increasing length and SNR, regardless of the size of the estimation window (Figure 4A. *Top*). In addition, when uncertainty distributions were plotted as a function of expected SMSS and ODC values, estimation of ODC was observed to be more difficult for extreme ODC input values and easier for higher input SMSS values (Figure 4B, *Left*) ^57–59^. Conversely, while SMSS uncertainty converged to zero with trajectory length, it appeared to depend very little upon the local SNR surrounding individual detections (Figure 4A, *Bottom*). In addition, the SMSS estimation was observed to be easier at the extremes of both input SMSS and ODC (Figure 4B, *Right*).

### OMEGA implementation of the method used to estimate the motion type classification error

The method to quantify the uncertainty associated with ODC and SMSS estimation described here has been implemented as part of our integrated OMEGA particle tracking and motion analysis data management software platform ^7^, with the specific goal of making it widely available to bench scientists regardless of their computational expertise (Figure 5 and Supplemental Figure 4). To this aim, OMEGA automatically keeps track of all data and metadata elements associated with the SPT, SNR estimation and motion analysis pipelines, making the entire process easy to execute and transparent to the user (Supplemental Figure 4).

**Figure 5:**
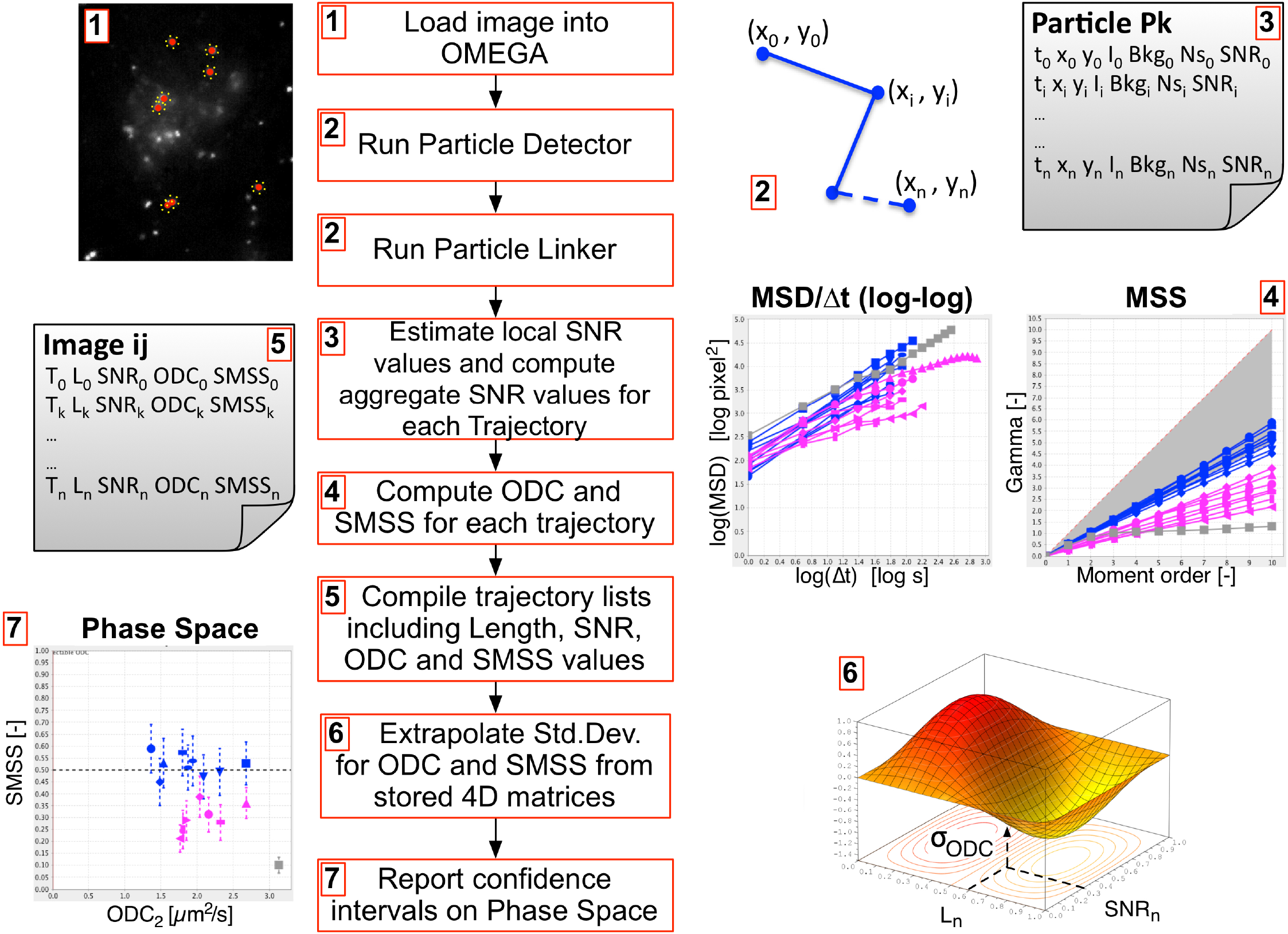
Workflow for the estimation and reporting of motion type estimation uncertainty associated with specific particle tracking and motion analysis runs in OMEGA. The method developed to compute and report the error associated with motion type measures computed in OMEGA entails the following steps: **(1)** Images are loaded onto OMEGA. **(2)** A given pair of particle detection/localization and linking algorithms, are used to estimate particle positions at each time point and link together successive positions of the same particle across time. **(3)** The position of individual particles is fed to the OMEGA SNR Estimation plugin to determine local Background (*Bkg_n_*), Noise (*Ns_n_*) and SNR (*SNR_n_*) values associated with each localized spot and compute representative aggregate SNR values for each trajectory (i.e., minimum local SNR per trajectory; TMinSNR). **(4)** ODC and SMSS estimations are computed for each trajectory. **(5)** For each particle tracking run, trajectory lists are compiled including trajectory ID (T_n_), trajectory length (*L_n_*), TMinSNR (*SNR_n_*), ODC (*ODC_n_*) and SMSS (*SMSS_n_*). **(6)** For each trajectory, this tuple of indices is utilized to interrogate pre-computed fourdimensional (4D) matrices (Figure 3, Step 11) and extrapolate the corresponding values of standard deviation for ODC and SMSS. **(7)** Obtained standard deviation values are utilized to compute uncertainty intervals on the basis of the desired confidence level (i.e., 2x *σ* corresponds to a 95% confidence) that are plotted on the ODC vs. SMSS phase space to indicate the confidence of motion type estimation and are reported in tabular form.

In order to quantify motion type estimation uncertainty in the context of OMEGA, selected images are retrieved from available image repositories (i.e., OMERO), loaded into OMEGA and subjected to particle detection using one of the supported algorithm (Figure 5). Detected particle coordinates are passed to the OMEGA SNR estimation plugin together with the corresponding image-planes, to obtain the P_PI_, local noise, local background and local SNR associated with each detected feature point. After linking, global ODC and SMSS estimates are computed for each homogeneous trajectory or segment using the OMEGA DTM plugin, and tables relating Particle Detection/Localization Algorithm ID and Trajectory ID with observed trajectory length (i.e., L_output_), minimum observed local SNR for all particles in the trajectory (i.e., T_MinSNR_), observed ODC (i.e., ODC_output_) and observed SMSS (i.e., SMSS_output_) are generated as follows (Figure 5):

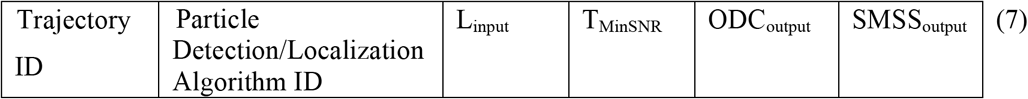

Upon user request data, expected ODC and SMSS estimation uncertainty values associated with each found trajectory (Figures 5 and 6) can be automatically extracted by linear interpolation from the four-dimensional data structures in Eq. 6 (Figure 3), on the basis of set of observed L, local SNR, ODC and SMSS trajectory parameters stored as in Eq. 7 (Supplemental Figure 4B). Once obtained, resulting uncertainty values are then reported in tabular form and used to compute confidence intervals (e.g., 2x *σ*corresponds to a 95% confidence interval) that are displayed on the two-dimensional ODC vs. SMSS phase space (Figure 6B and E), which represents the keystone of the 4-plots (i.e., x vs. y coordinates; MSD vs. t log-log; MSS; and D vs. SMSS phase space plots) motion type classification method ^7,40^ implemented in OMEGA (Figures 5-7, 6C and 6D).

**Figure 6:**
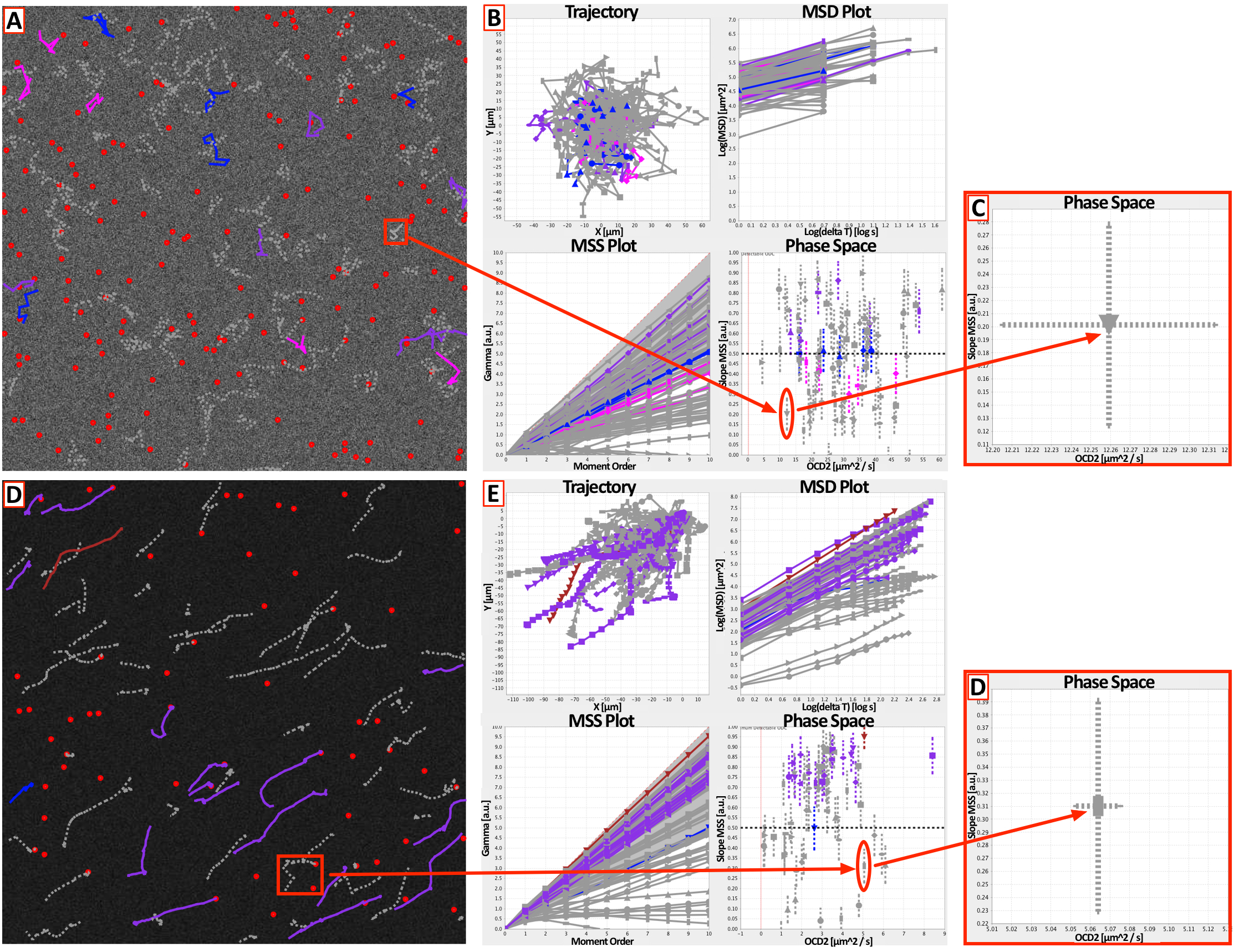
OMEGA diffusivity measures uncertainty estimation use-case using standardized multiple particle tracking benchmarking datasets mimicking viral particle movement in infected cells. Benchmarking time series from the Chenouard *et al.*, multiple particle tracking benchmarking dataset (Chenouard *et al.*, 2014) corresponding to mobility scenario IV (virus), low particle density, and SNR = 1 (A) vs. SNR = 7 (D), were subjected to SPT and motion analysis in OMEGA. Images were loaded onto OMERO and imported into OMEGA using the OMEGA image browser. Images were subjected to single particle tracking using the OMEGA implementations of the MOSAICsuite Feature Point Detection and Linking algorithms (Sbalzarini & Koumoutsakos, 2005). (**A and D**) Resulting particles and trajectories were overlaid over the corresponding image using the OMEGA side bar image viewer. Trajectories that displayed a straight Moment Scaling Spectrum plot (Ferrari *et al.*, 2001) indicating uniform behavior for the duration of motion, were assigned a color-code corresponding to their apparent motion type using the OMEGA Trajectory Segmentation (TS) plugin and color code conventions (*fuchsia*, sub-diffusive; *purple*, super-diffusive; *blue*, Brownian). (**B and E**) All trajectories were subjected to diffusivity analysis using the OMEGA Diffusivity Tracking Measure (DTM) plugin and plotted on the 4-plots set (xy, MSD vs. t log-log, MSS and D vs. SMSS phase space plots) method used for motion type classification in OMEGA (Ewers *et al.*, 2005). (**C and F**) The phase-space behavior and corresponding error bars of two representative trajectories of similar length selected from the two SNR scenarios are displayed in the zoomed in inset as indicated.

### Example use case of ODC and SMSS uncertainty estimation

In order to provide an example use case for the method described here, standardized benchmarking time series from the Chenouard et al. multiple particle tracking benchmarking dataset ^32^ were subjected to the SPT, motion analysis and error estimation workflow in OMEGA (Figure 6) ^7^. Times series mimicking viral particle mobility (i.e., mobility scenario IV - VIRUS), with low particle density, and with two representative SNR conditions (i.e., SNR = 1 vs. SNR = 7) were loaded onto OMERO and imported into OMEGA using the OMEGA Image Browser. Images were subjected to single particle tracking using the OMEGA implementations of the MOSAICsuite FPD and Linking algorithms ^49,55^. Resulting particles and trajectories were then overlaid over the corresponding images using the OMEGA sidebar image viewer (Figure 6A and D). Subsequently, trajectories that displayed a straight Moment Scaling Spectrum (MSS) plot indicating uniform behavior for the duration of motion ^40,48^, were assigned a color-code corresponding to their apparent motion type, using the OMEGA Trajectory Segmentation (TS) plugin and following the color-convention used in OMEGA ^7^. Thus, trajectories with straight MSS lines and slope below 0.5 were considered sub-diffusive and assigned the color *fuchsia*, trajectories with MSS slope > 0.5 were considered super-diffusive and assigned the *purple* color, and trajectories with MSS slope ~ 0.5 were assigned the color *blue*, corresponding to Brownian motion. Finally, all trajectories were subjected to diffusivity analysis using the OMEGA DTM plugin and plotted using the 4-plots method used for motion type classification in OMEGA (Figure 6B and E) ^7,40^. While, with benchmarking images synthesized to reflect higher SNR (Figure 6D) most resulting trajectories displayed the expected directionally biased behavior mimicking Levy flights ^60^, in the situation with lower SNR conditions (Figure 6A) trajectories appeared less “stretched out”. Consistently, most trajectories from images with SNR = 7 scenario clustered in the top half of the phase space graph (Figure 6E), with a prevalence of SMSS values >> 0.5, indicating super-diffusive or directed behavior. Conversely, trajectories obtained from the SNR = 1 scenario displayed a markedly reduced apparent directionality, with several trajectories displaying SMMS close to 0.5, indicating a pseudo-diffusive behavior. Because, the mobility behavior for both image sets was set to mimic viral dynamics independently of the simulated SNR conditions ^32^, one possible explanation of this discrepancy was the higher frequency of localization and linking errors associated with lower image quality. This difference was underscored by the markedly higher uncertainty levels for ODC estimation observed with lower vs. higher SNR values, which is also consistent with the stark dependency of ODC uncertainty on SNR observed in Figure 4. Conversely, the error bar for SMSS appeared to be similar in both SNR scenarios, which is similarly consistent with what observed in Figure 4, where the uncertainty of SMSS appears to be much more dependent on length than on SNR.

## Materials and Methods

### Benchmarking test cases

In order to construct appropriate test cases for simulation and validation purposes, the input values of P_PI_, and of trajectory length (L), ODC and SMSS were set as described in Supplemental Table I. Unless otherwise indicated, all test cases datasets contained 1000 independently generated images or trajectories as appropriate. In all cases, validations were performed assuming the following physical dimensions: 1 pixel/μm and 1 frame/sec.

### Algorithms description and validation

#### Artificial image generation and validation

Artificial images were generated following a described method ^61^ using a Java program called, *artificialImageGenerator* which was developed ^7^ to reproduce a previously described MatLab routine ^49^. The functionality of the Java program was evaluated by direct comparison with the MatLab version of the algorithm (Supplemental Figure 1). In brief, sets each containing 1000 independently generated 118 x 118 pixel synthetic planes representing moving blobs were created to contain ten simulated point sources moving on an horizontal straight line at a constant pixel displacement per frame (i.e., 0.27 pixel/frame) and were initialized by adding a background (black) intensity value of B = 10 to each pixel. In order to model different image qualities (vis. SNR), point sources were assigned 12 increasing P_PI_ values (Supplemental Table I) and microscopic observation was simulated by sampling a Gaussian of g = 1 pixel and μ = P_PI_, centered at each known point position. In order to model Poisson-distributed Shot noise associated with CCD camera image acquisition, the peak intensity P_PI_ of each pixel was replaced with a number sampled at random from a Poisson distribution of mean value μ = P_PI_. All random numbers were generated independently for every trial and plane. Artificial planes are stored as unscaled 16-bit TIFF files. For validation purposes, these planes were directly compared (i.e., pixel-per-pixel) to those generated using the published algorithm ^49^. As shown in Supplemental Figure 1, discrepancies measured either as the number of discordant pixels (Supplemental Figure 1, Top) or as the distance from ground-truth (DGT; Supplemental Figure 1, Bottom) of the observed mean intensities over the entire image, were very rare and consistent to numerical errors due to well known rounding differences between MatLab and Java and possibly due to the use of different hardware.

#### Generation of distributions of particle detection/localization errors

Twelve test sets each characterized by an increasing value of input particle peak intensity P_PI_ (Supplemental Table I), and each containing 1000 independently generated 20 x 20 pixel artificial image planes were produced using the Java *artificialImageGenerator* program ^7,49^, with the exception that individual simulated point sources placed at random known sub-pixel positions (i.e., ground truth) within the central pixel of the plane were used instead of moving particles.

The stationary feature points present in each synthetic plane were then localized using the OMEGA MOSAICsuite FPD plugin ^49,55,56^, and the input Cartesian position of each simulated source points were compared with their observed position. Distributions of observed distance from ground truth (DGT) of the Cartesian PosV (i.e., DGT_PosV_) were computed and sampled uniformly at random to obtain pairs of coupled particle position offsets values.

#### Generation and validation of artificial trajectories of known input ODC and SMSS

A total of 1936 test sets each containing 1000 independently generated artificial trajectories of arbitrary motion type and known 16 L x 11 ODC x 11 SMSS input values (Supplemental Table I), were generated using the *artificialTrajectories2* Matlab routine developed earlier for this purpose ^7,62^ and used as a basis for “noisy” trajectory generation (see below).

Because of the inherently stochastic nature of any motion process prevents point-by-point trajectory comparisons, the only available avenue to validate the quality of artificial trajectories is to contrast trajectory measures obtained from a test vs. a control set. Thus for validation purposes, artificial trajectories generated using *artificialTrajectories2* were compared with equivalent Brownian trajectories of known L and ODC (Supplemental Figure 5) generated by employing our *brownian1* MatLab routine ^7^. Specifically, our previously developed *getMSS* Matlab function ^7,59,62^ was used to back compute SMSS from benchmarking sets of 1000 simulated trajectories generated with *artificialTrajectories2* to reflect 17 L x 11 ODC x 1 SMSS (SMSS = 0.5) input values (Supplemental Table I; Supplemental Figure 5A) and the results obtained with L/3, L/5 and L/10 calculation-window sizes were compared with those obtained from equivalent sets of Brownian trajectories. It should be noted here that because *artificialTrajectories2* is internally dependent on *getMSS* to generate trajectories of known ODC, the analysis was restricted to comparisons of input vs. output values of SMSS. As shown, when the absolute values of the relative error of SMSS (|Relative DGT_SMSS_|) were computed and plotted for artificial trajectories generated using *artificialTrajectories2* vs. *brownian1*, results obtained were very similar to each other regardless of L or input D even though random-walks displayed in general a higher variance across all conditions (Supplemental Figure 5A).

In order to extend the validation to a more complete range of test-cases, 1870 test sets, each containing 1000 artificial trajectories (Supplemental Table I), were produced using *artificialTrajectory2* and SMSS values were back computed using the OMEGA DTM plugin as above ^7^. When observed DGT_SMSS_ distributions were plotted as a function of calculation-window, input L, ODC and SMSS values (Supplemental Figure 5B), we observed that their bias and sigma were largely independent from input ODC while they rapidly converged to 0 with increasing input L and SMSS. As expected, DGT_SMSS_ was observed to be strongly depended on the size of the calculation-window with variance significantly decreasing with increasing windows sizes.

#### Production and validation of “noisy” trajectories

“Noisy” trajectories were produced using the *noisyTrajectoryGeneration* MatLab routine developed for this study. Briefly, using this routine the effect of detection/localization error on the 1936 test sets of infinitely precise trajectories generated above, was modeled by replacing each point, P_i_ = (x_i_, y_i_) along a given trajectory with a new “noisy” point, noisy_P_i_ = (noisy_x_i_, noisy_y_i_), whose position was obtained as follows:

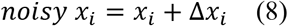

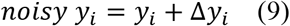

Where, each value was independently obtained by sampling uniformly at random distributions of paired DGT_PosV_ = offsets, obtained for the same set of 12 local SNR_input_ values as described above (Figure 3; Supplemental Figures 2 and 3)

The validation of “noisy” trajectories, was carried out by comparing input with output particle position offsets (Supplemental Figure 6) as follows. The effect of localization uncertainty on artificial trajectories was modeled for three artificial trajectory test sets each containing 1000 infinitely precise trajectories of known motion type characteristics and length (L = 50, 250 and 1000; D = 0.01 and SMSS = 0.5) and mimicking the effect of four peak intensity values (I = 15.00, 28.73, 154.70, 1000.00). Trajectories were first generated using our *artificialTrajectories2* MatLab function and then detection uncertainty was modeled for each of the four input peak intensity values by adding to each point coordinate pairs of Δx and Δy offsets (i.e., Input offsets) sampled uniformly at random from previously computed distributions (Supplemental Figure 6). Subsequently, offsets were back computed (i.e., *Output* offsets) and the distributions of Input vs. Output offsets for the x coordinate were compared directly (Supplemental Figure 6A). In addition, the distance between Input and Output offsets distributions for both coordinates were quantified by using the Wasserstein Distance (WD) metric (Supplemental Figure 6B) ^63^. Artificial distributions with a WD metric equal to 0.00, 1.47 and 3.00 respectively are displayed for comparison sake (Supplemental Figure 6C). While Input and Output offsets distributions showed excellent overlap in all test cases, we observed that similarity nonetheless increased with both peak intensity and trajectory length which is expected as error is known to be strongly dependent on image quality and sampling size.

## Discussion

Known mathematical properties of a specific measurement or calculation can be used to <u>theoretically predict</u> the amount of uncertainty associated with it. Conversely, error can be <u>estimated empirically</u> from the behavior of independently repeated measurements or calculations of the same quantity. Attempts to theoretically predict the error associated with ODC and MSD calculations have been made ^64,65^. While Martin’s error model ^64^ was initially defined for MSD, it was subsequently extended to moments of displacement of higher order allowing its theoretical application to SMSS computations ^57,58^. However because the validity of the model in these new conditions was not established and tracking errors remain difficult to theoretically predict, it was reasoned that a better approach would be to empirically estimate the uncertainty associated with each ODC and SMSS measurement (i.e., local error analysis). This is further underscored by the conclusions reached by Michalet and Berglund ^65^ that localization imprecision fundamentally limits the accuracy with which ODC can be estimated especially during frequent motion type switching regimes, such as are encountered in several commonly encountered experimental scenarios. In this context, the empirical quantitation of uncertainty is of paramount importance as it has the potential of strongly restraining scientific conclusions that can be drawn from experimental data.

The main sources of error in SPT are ascribable to particle detection and localization, and to linking ^66,67^. When attempting to localize a point emitter from a finite number of detected photons, an estimate of the true spatial location may be obtained using a specific detection/localization algorithm ^68^. Background autofluorescence, motion blur, finite photon count, camera pixel resolution, and camera type-specific noise all contribute to the inaccuracies of each specific algorithm and cause the estimated position to differ from the true molecular position by an error, ε, that is equal to the difference between the observed and the ground truth position. This distance from ground truth (DGT) is typically assumed to be a random variable whose standard deviation, *σ*, measures the indeterminate component of the error (i.e., localization precision reflecting the closeness of agreement between independent positional estimates, and generally resulting from imaging fluctuations and noise), and whose mean, *μ*, measures the determinate component of the error (i.e., closeness between the mean of observed positions and the true position and generally dependent upon algorithm accuracy). This study analyzes the dependence of the accuracy and precision of particle localization on the duration of movement (i.e., the length of the trajectory) and the SNR of the images, which both strongly affect the uncertainty with which the movement of single particles can be analyzed and trajectories can be classified on the basis of their observed motion characteristics ^64,69^. However, a limitation of this method is that it does only partially consider motion type estimation errors that are due to linking mistakes. Linking errors can have two effects: (1) If links are mistakenly not detected, trajectories will appear shorter than the ground truth, thus increasing sampling errors. (2) If spurious links are detected within a trajectory, motion characteristics can be biased, thus indirectly affecting estimation uncertainty. While our method considers the former, it does not consider the latter. Errors due to wrong links are typically stark (outliers) and can be identified by statistical methods (outlier detection) or sanity checks on the results. Nonetheless, the errors presented here should be considered as lower bounds and the real error might be higher.

Even with infinitely accurate and precise positioning, global trajectory measures are nonetheless expected to display statistical variance because of sampling errors (i.e., finite trajectory lengths), which diminishes as the number of points that are available for calculation. On this basis, localization error inevitably enhances the uncertainty with which measures such as ODC and SMSS, can be estimated. Furthermore, in addition to positional uncertainty and sampling, it has been predicted that the observed dynamics of the sub-diffractive moving object also affects the accuracy of ODC and SMSS estimation ^57,58^. In particular, the variance of the *ρ*-th displacement moment depends on both *ρ* and on the value of the ODC of order *ρ*, thus also affecting SMSS ^57–59^. Consistently, results presented here and obtained with the OMEGA Diffusivity Tracking Measures plugin ^7,40^, indicate that the accuracy of ODC estimation increases with trajectory length and SNR and starkly depends upon observed ODC and SMSS values of individual trajectories (Figure 4). Conversely, while in our hand SMSS estimation accuracy increased with length and was clearly dependent upon both input SMSS and ODC, it displayed a much lower dependency upon noise level and associated particle localization imprecision than predicted underscoring the importance of performing simulation studies to substantiate theoretical claims.

## Conclusions

Because localization precision is intimately dependent upon the noise distribution and the SNR of the images, the first step of the uncertainty estimation method described here consists of obtaining the error distribution associated with a specific algorithm one wishes to use for particle tracking and with a representative range of SNR values. Having established the accuracy and precision of each algorithm under preset SNR conditions, the next step is to empirically assess how such localization uncertainties affect the quality of ODC and SMSS estimates. In order to do so, simulated trajectories of known length, particle positions and motion type characteristics (i.e., known ODC and SMSS values) are produced, and distributions of paired x and y coordinate errors associated with each input SNR value are sampled uniformly at random to offset the position of all points along each trajectory. These “noisy” trajectories are then used to back compute ODC and SMSS in order to extract distributions of distance from ground truth for both measurements, calculate their standard deviations and store them for each combination of input L, ODC, SMSS and SNR parameters. In the final step of the method, these stored matrices are interrogated via interpolation each time particle tracking and diffusivity analysis is performed to associate each value of ODC and SMSS with its predicted uncertainty.

A comprehensive understanding of any complex biological process, such as for example viral infection, requires real-time interrogation of the dynamic behavior of multiple molecular components across a wide range of biological model systems, sample types, experimental perturbations, and spatiotemporal scales. In the face of this inherent multidimensionality and complexity, scientific reproducibility can only be guaranteed by the use of integrated platforms capable of automatically account for data provenance, quality control, multiple data types and techniques, and last but not least the standardized and accurate evaluation and reporting of the propagation of error through the analytical pipeline. This is particularly relevant in biomedical imaging, which often requires the quantitative analysis of gigabytes of multi-dimensional, feature-rich, high-resolution digital images as a prerequisite for understanding the molecular underpinnings of biology and disease. Because of the inherent heterogeneity of most subcellular processes, high spatiotemporal-resolution imaging of the realtime dynamic behavior of individual intracellular macromolecules, molecular complexes and organelles, coupled with singleparticle tracking (SPT), and motion analysis is increasingly used to extract quantitative parameters on single molecules and their environment and to discern their biological function.

Interpreting images as scientific measurements, rather than as mere qualitative observations, requires the development of shared methods for uncertainty quantification and error analysis. This is particularly true in case of large collaborative projects that produce large amounts of data and stores only the final analysis result, where uncertainty quantification combined with the use of collaborative software and re-usable algorithms is critical for data interpretation, reproducibility and dissemination, and for efficient problem solving and processing. The algorithm-centric assessment of how noise in the raw image affects uncertainty in the analysis results can be addressed by analyzing the error-propagation behavior of a given algorithm. While this approach is commonplace in numerical analysis and applied mathematics ^52,54^, to our knowledge our work represents the first attempt of performing such an analysis for uncertainty quantification in particle-tracking trajectory analysis.

Work presented here addresses the question of how accurately rate of displacement, as measured by the observed diffusion constant (ODC) ^5,57^, and freedom of motion, as measured by the Hurst exponent, also known as the slope of the moment scaling spectrum or SMSS ^40,47,49,59^, can be estimated from the analysis of finite length single-particle trajectories whose individual points are significantly affected by localization errors. Because this question is of paramount importance for modern biology, the ultimate goal is to propose a method that could become the basis for a standard in the assessment of motion type estimation uncertainty. For this purpose the novel algorithm-centric Monte Carlo simulation method to empirically assess and report motion type estimation error described here, was designed to be easily adaptable to different noise models and particle-tracking algorithm. In addition, to facilitate dissemination and standardization it was incorporated in our recently released Open Microscopy Environment inteGrated Analysis (OMEGA) software platform ^7^ and described in our Minimum Information About Particle Tracking Experiments (MIAPTE) metadata standard ^8^. We hope this example will be applied to other image analysis problems and will serve as a basis for the standardized development of generic methods for uncertainty quantification in biomedical imaging.

## Acknowledgements

The success of this multidisciplinary project was due to the collective effort of many individuals. We want to specifically acknowledge our debt of gratitude toward those without whom this software would simply not have been produced. We are deeply beholden to **Mario Valle** (CSCS - Swiss National Supercomputing Centre, ETH-Zurich), **Peter Kunszt** (SystemsX.ch now at Dynatrace Barcelona) for continual encouragement, critical support (financial and otherwise), and lots of helpful discussions. <u>Without Mario and Peter this project would never have started!</u>

We thank **Raffaello Giulietti** and **Roberto Mastropietro** (University of Applied Sciences and Arts of Southern Switzerland - SUPSI), for their assistance and guidance, for invaluable discussions, and for their generosity with their software development and project management expertise during pivotal times along our path. We are extremely grateful to **Karl Bellvé, Kevin Fogarty** and **Lawrence Lifshitz** (University of Massachusetts Medical School) for their support and encouragement, their generosity with code, time, knowledge and experience, and for critical discussions and feedback. We are indebted to **Kevin Eliceiri** (LOCI, University of Wisconsin at Madison), **David Grunwald**, Lawrence Lifshitz and **Thoru Pederson** (University of Massachusetts Medical School, Worcester, MA), and **Orlando Petrini** (Istituto Cantonale di Microbiologia, Ticino now at POLE Pharma Consulting), for stimulating discussions, inspiration and motivation. We are immensely grateful to **Diego Frei, Raffaello Giulietti, Loris Grossi**, and **Tiziano Leidi** (University of Applied Sciences and Arts of Southern Switzerland - SUPSI), **Pietro Incardona**, (Max-Plank Institute of Molecular Cell Biology and Genetics - Dresden), and Lawrence Lifshitz, for sharing their code and software engineering skills with A.R. We thank the entire **Open Microscopy Community** (openmicroscopy.org) for inspiration, encouragement, motivation and help. They have been our role models throughout and as such their contribution is simply too extensive to be properly documented. Our gratitude extends but is not limited to: **Chris Allan, Sebastian Besson, Jean-Marie Burel, Melissa Linkert, Josh Moore, Will Moore**, and **Jason Swedlow** (OME and Glencoe Software), **Christian Dietz**, Kevin Eliceiri and **Curtis Rueden** (ImageJ/Fiji and KNIME ecosystem). We are grateful to Mario Valle for critically reviewing the manuscript. We apologize to colleagues whose work we were not able to cite due to space limitations.

Extramural funding was from the Swiss National Science Foundation (Project CRSII3_136282 to C.-S.-D.-C.), and the European Commission FP7 (Project HEALTH-2007-2.3.2, GA HEALTH-F3-2008-201,032, to C.-S.-D.- C.). Institutional funding was from the **Institute of Research in Biomedicine**, the **University of Geneva** (to C.-S.-D.-C.), **SystemsX.ch Information Technology, Emory University** (to A.R., and V.G.), **University of Applied Sciences and Arts of Southern Switzerland - SUPSI** (to A.R., and V.G.), and the **University of Massachusetts Medical** school (to A.R., C.-S.-D.-C., and V.G).

## Supplemental Material

**Supplemental Figure 1: Validation of artificial image generator.**

**Supplemental Figure 2: Simulated single point emitters were used to empirically estimate detection and localization uncertainty.**

**Supplemental Figure 3: Distribution of particle localization error observed with the particle detection/localization algorithm implemented in OMEGA.**

**Supplemental Figure 4: Motion type estimation uncertainty quantitation data flow as implemented in OMEGA.**

**Supplemental Figure 5: Validation of procedure for artificial trajectories generation.**

**Supplemental Figure 6: Validation of procedure used to generate “noisy” artificial trajectories.**

**Supplemental Table I: Benchmarking test cases.**

## Abbreviations

DGT: distance from ground truth
MIAPTE: minimum information about particle tracking experiments
MPT: multiple particle tracking
MSD: mean squared displacement
MSS: moment scaling spectrum
ODC: observed diffusion constant
OMERO: open microscopy environment remote objects
SNR: signal to noise ratio
TS: Trajectory Segmentation

